# SARS-CoV-2 Delta derivatives impact on neutralization of Covishield recipient sera

**DOI:** 10.1101/2021.11.30.470521

**Authors:** Rima R. Sahay, Deepak Y. Patil, Gajanan N. Sapkal, Gururaj R. Deshpande, Anita M. Shete, Dimpal A. Nyayanit, Sanjay Kumar, Shanta Dutta, Pragya D. Yadav

## Abstract

The emergence of SARS-CoV-2 Delta variant and its derivatives has created grave public health problem worldwide. The high transmissibility associated with this variant has led to daily increase in the number of SARS-CoV-2 infections. Delta variant has slowly dominated the other variants of concern. Subsequently, Delta has further mutated to Delta AY.1 to Delta AY.126. Of these, Delta AY.1 has been reported from several countries including India and considered to be highly infectious and probable escape mutant. Considering the possible immune escape, we had already evaluated the efficacy of the BBV152 against Delta and Delta AY.1 variants. Here, we have evaluated the neutralizing potential of sera of COVID-19 naive vaccinees (CNV) immunized with two doses of vaccine, COVID-19 recovered cases immunized with two doses of vaccine (CRV) and breakthrough infections (BTI) post immunization with two doses of vaccine against Delta, Delta AY.1 and B.1.617.3 using 50% plaque reduction neutralization test (PRNT50). Our study observed low NAb titer in CNV group against all the variants compared to CRV and BTI groups. Delta variant has shown highest reduction of 27.3-fold in NAb titer among CNV group compared to other groups and variants. Anti-S1-RBD IgG immune response among all the groups was also substantiated with NAb response. Compromised neutralization was observed against Delta and Delta AY.1 compared B.1 in all three groups. However, it provided protection against severity of the disease and fatality.

## Text

Over the course of Coronavirus disease (COVID-19) pandemic, several factors had an impact on increasing Severe acute respiratory syndrome Coronavirus −2 (SARS-CoV-2) infections. These factors include human behavior, inappropriate infection prevention practices, waning or poor immune response after natural infection or post vaccination. The emergence of SARS-CoV-2 variants of concerns (VOCs) with distinctive mutations made the pandemic situation gruesome. A devastating second wave emerged globally fueled by the highly transmissible and increased pathogenic Delta variant. This VOC was first identified in India and subsequently reported from at least 147 countries worldwide.^1^ In a short span of time Delta variant has rapidly mutated into several sub-lineages known as ‘Delta derivates’ (Delta AY.1 to AY.126).^2^ Of these, AY4.2 is currently increasingly being reported in United Kingdom and also detected in Europe, including Denmark, Germany and Poland.^2^ This was preceded in the past with, another Delta sub-lineage Delta AY.1 and was said to be more threatening than the Delta variant. Delta AY.1 was initially identified in India and traced to many cases across the globe^3^ and has raised a concern about the vaccine efficacy. AY.1 variant have ≥20% high prevalent mutation than the Delta variant which has T19R, E156G, W258L, K417N, L452R, T478K, D614G, P681R and D950N amino acid mutations in the spike gene. Although the worldwide cumulative prevalence of Delta AY.1 (<0.5%) is lower than Delta variant (17%), it is still not clear whether Delta AY.1 variant is deadlier than Delta.^1,3^ Delta variant is the major reason for the breakthrough infections world over.^4^ Several studies have reported the neutralization potential of the approved COVID-19 vaccines against Delta variant.^4^ Studies have also observed 4 to 9-fold reduction in the neutralization antibody (NAb) titer with sera of ChAdOx1 vaccinees against Delta variant.^4^ Liu et al., have reported the susceptibility of Delta AY.1 variants to neutralization with the sera of BNT162b2 vaccinee.^5^ We have previously demonstrated minor reduction in the NAb titer of the sera of COVID-19 recovered individuals with BBV152 vaccination and breakthrough cases against Delta AY.1 compared to Delta variant.^6^ However, the data on the neutralization potential of the approved COVID-19 vaccines against Delta AY.1 still remains limited. Considering this, we have assessed the susceptibility of Delta, Delta AY.1 and B.1.617.3 variants to Covishield vaccine (ChAdOx1 produced by Serum Institute of India Limited) which is widely administered in India.

We determined the NAb titre of sera obtained from 25 COVID-19 naïve vaccinees (CNV) immunized with two doses of vaccine (8 weeks post vaccination), 16 COVID-19 recovered cases immunized with two doses of vaccine (CRV) (8 weeks post vaccination) and 26 breakthrough infections (BTI) post immunization with two doses of vaccine against Delta, Delta AY.1 and B.1.617.3 using 50% plaque reduction neutralization test (PRNT50). Prototype B.1 variant was used for comparative analysis.

## Methods

Briefly, the sera of study subjects were serially four-fold diluted and further mixed with an equal amount of Delta, Delta AY.1, B.1.617.3 and B.1 virus suspension (50-60 Plaque forming units in 0.1 ml) separately. Post the incubation at 37°C for 1 hr, each virus-diluted serum sample (0.1 ml) was inoculated onto duplicate wells of a 24-well tissue culture plate of Vero CCL-81 cells. After incubating the plate at 60 min, an overlay medium consisting of 2% Carboxymethyl cellulose (CMC) with 2% fetal calf serum (FCS) in 2× MEM was added to the cell monolayer after incubation of the plate at 37°C for 1 hr. The plate was further incubated at 37°C with 5% CO2 for 5 days. At assay termination, plates were stained with 1% amido black for an hour. Neutralizing antibody titers were defined as the highest serum dilution that resulted in >50 (PRNT50) reduction in the number of plaques.

An IgG immune response was also assessed using anti-S1-RBD ELISA.^7^ Briefly, 96-well ELISA plates (Nunc, Germany) were coated with SARS-CoV-2 specific antigen (S1-RBD at a concentration of 1.5μg/well in PBS pH 7.4). The plates were blocked with a Liquid Plate Sealer (CANDOR Bioscience GmbH, Germany) and Stabilcoat (Surmodics) for two hours at 37°C. The plates were washed two times with 10 mM PBS, pH 7.4 with 0.1 per cent Tween-20 (PBST) (Sigma-Aldrich, USA). The sera of the subject from CNV, CRV and BTI group (1:50 dilution) were serially four-fold diluted and added to the antigen-coated plates and incubated at 37°C for one hour. These wells were washed five times using 1× PBST and followed by addition 50 μl/well of anti-human IgG horseradish peroxidase (HRP) (Sigma) diluted in Stabilzyme Noble (Surmodics). The plates were incubated for half an hour at 37°C and then washed as described above. Further, 100 μl of TMB substrate was added and incubated for 10 min. The reaction was stopped by 1 N H2SO4, and the absorbance values were measured at 450 nm using an ELISA reader. Anti-SARS-CoV-2 antibody (NIBSC code 20/130) was also included in the assay as a reference standard. The cut-off for the assay was set at twice of average OD value of negative control. The endpoint titer of a sample is defined as the reciprocal of the highest dilution that has a reading above the cutoff value. The statistical significance was assessed using a two-tailed Kruskal-Wallis test with Dunn’s test of multiple comparisons. P-value less than 0.05 were considered to be statistically significant for the applied test.

## Results and Discussion

The geometric mean titer (GMT) of NAb against B.1, Delta, Delta AY.1 and B.1.617.3 were determined with the sera of the subjects from CNV group (30.8, 1.1, 9.8, 1.6), CRV group (1248, 489.8, 403.8, 314) and BTI group (876.7, 499.8, 415.8, 235.8) respectively (Figure 1). The magnitude of anti-S1-RBD antibody immune response among CNV group (range 100-3200, mean 1312), CRV group (range 1000-3200, mean 2717) and BTI group (range 400-3200, mean 1933) correlated with neutralization. NAb titer in CNV group was found to be very low against all the variants compared to CRV and BTI groups. This suggests the need for the booster vaccination to cope up with waning immune response among vaccinees. Delta variant has shown highest reduction of 27.3-fold in NAb titer among CNV group compared to other groups and variants. This finding implies highest probability of the breakthrough infection with Delta variant. Of the 26 cases from BTI group, SARS-CoV-2 genome sequences of 14 breakthrough cases could be retrieved using Next generation sequencing. These cases were found to be affected with Delta (n=8) and Kappa (n=6) variants. A total of 21 cases were found to be asymptomatic; while 5 cases had mild disease. Reduction in the NAb titer was also observed in CNV (19.17-fold), CRV (3.97-fold), BTI (3.72-fold) groups with B.1.617.3 variant. This necessitates the exploration of vaccine efficacy of currently available COVID-19 vaccines against B.1.617.3 and other variants under monitoring.

**Figure 1:**
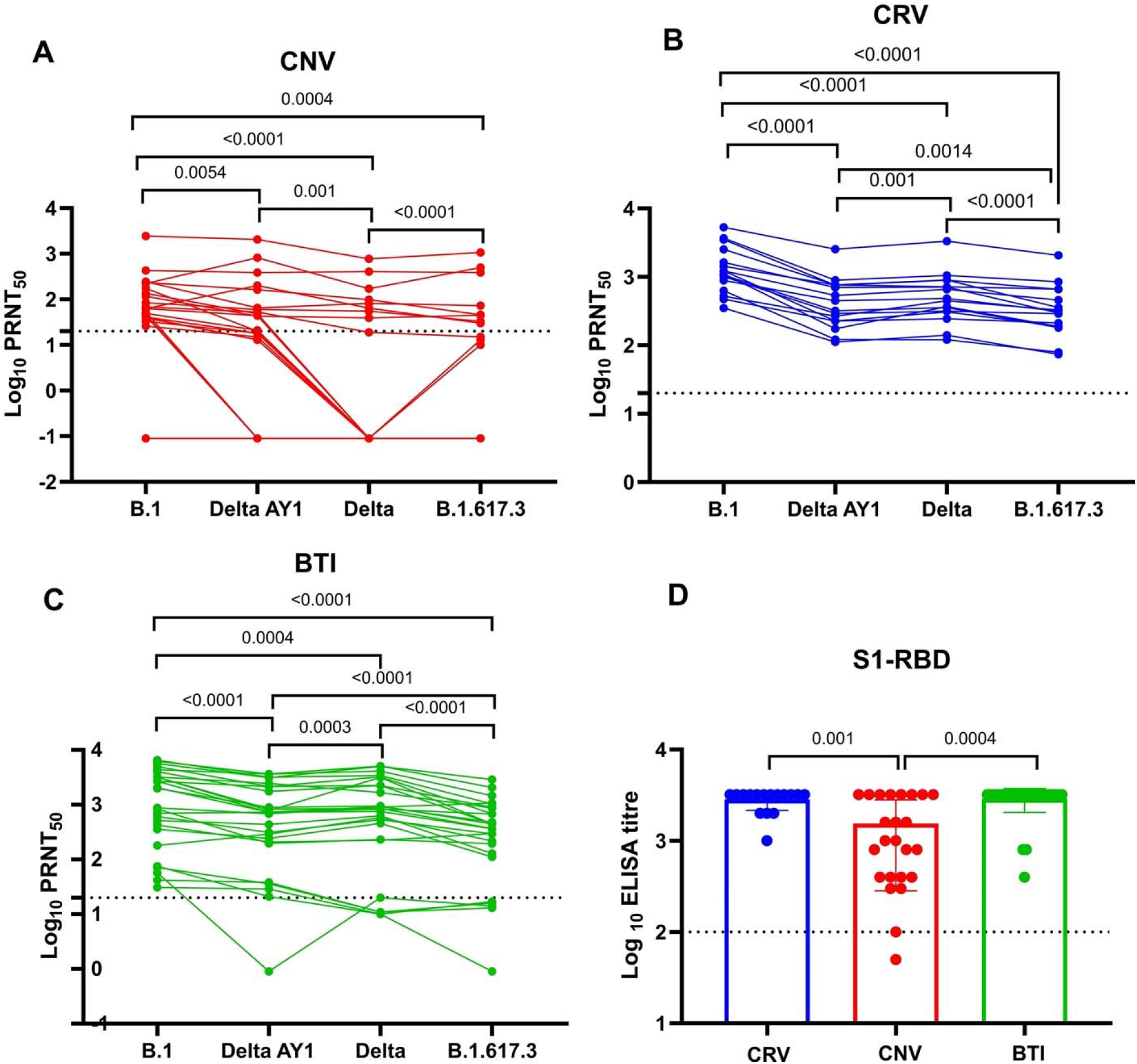
Neutralization of individual sera vaccinated with Covishield vaccine from different scenarios against B.1, Delta AY.1, Delta and B.1.617.3 strains and ELISA titer of individual sera vaccinated from different scenarios: Comparisons of neutralization titer of individual cases sera immunized with two dose vaccine Covishield (CNV), recovered case sera immunized with two dose vaccine Covishield (CRV) and breakthrough cases post two doses of Covishield vaccine (BTI). The B.1 (EPI_accession: EPL_ISL_825088) (a), delta AY1 (EPI_accession: EPI_ISL_2671901) (b), delta (EPI_accession: EPI_ISL_2400521) (c), and B.1.617.3 (EPI_accession: EPL_ISL_2497905) (d). The statistical significance for the neutralizing titer was assessed using a matched pair Wilcoxon signed-rank. Anti-SARS-CoV-2 IgG titers of CRV, CNV and BTI’s sera for inactivated SARS-CoV-2 (h), S1-RBD protein (i). The dotted line on the figures indicates the limit of detection of the assay. Data for both the neutralizing titer and IgG titers are presented as mean values +/- standard deviation (SD).

Several studies have reported the increase in immune response in COVID-19 recovered cases and breakthrough infections post vaccination.^4^ Such a rise in immune response effectively neutralizes the immune escape due to mutants among breakthrough cases compared to naïve COVID-19 cases. The main reason for waning immunity post vaccination in SARS-CoV-2 infected individuals or COVID-19 naïve vaccinees is short time survival of plasma blast cells.^8^ Once the individual get re-infection after vaccination or vaccination post recovery, memory B cells are triggered that generate higher level of immune response.^8–10^ In conclusion, our findings suggest that Covishield vaccine was able to neutralize Delta derivatives and prevent serious disease and fatality among breakthrough cases. A booster dose vaccination of COVID-19 naïve vaccinees would achieve protective immune response to fight against emerging SARS-CoV-2 variants.

## Ethical approval

The study was approved by the Institutional Human Ethics Committee of ICMR-NIV, Pune, India under the project ‘Assessment of immunological responses in breakthrough cases of SARS-CoV-2 in post-COVID-19 vaccinated group’.

## Financial support & sponsorship

The financial support was provided from Indian Council of Medical Research (ICMR), New Delhi under intramural funding ‘COVID-19 to ICMR-National Institute of Virology, Pune for conducting this study.

## Author Contributions

PDY contributed to study design, data analysis, interpretation and writing. RRS, DYP, GS, DAN, AMS and SK contributed to data collection, data analysis, interpretation and writing.

## Conflicts of Interest

Authors do not have a conflict of interest.

## Acknowledgement

Authors gratefully acknowledge the encouragement and support extended by Prof. (Dr.) Balram Bhargava, Secretary to the Government of India Department of Health Research, Ministry of Health & Family Welfare & Director-General, ICMR, New Delhi. We sincerely acknowledge the kind support of Prof. Priya Abraham, Director, ICMR-NIV, Pune. We sincerely acknowledge the excellent technical support of Ms. Aasha Salunkhe and Mr. Chetan Patil, during the study.

## Notes

### Competing Interest Statement

The authors have declared no competing interest.

